# Micro-mechanical approaches to characterize tip growth: Insights into Root Hair Elasto-Viscoplastic Properties

**DOI:** 10.1101/2025.08.06.668718

**Authors:** Thomas Alline, Léa Cascaro, David Pereira, Atef Asnacios

## Abstract

Root hairs are outgrowths of the epidermal cells of plant roots. They increase the root’s exchange surface with the soil and provide it with good anchorage in the soil. Root hairs are an emblematic model of apical growth, a process also used by yeasts and hyphae to invade their environment. From a mechanical perspective, the root hair is considered as an elastic cylinder under pressure, closed by a dome that behaves like a yield fluid. We introduce here two innovative mechanical setups and protocols to characterize the mechanical properties of single growing root hairs in Arabidopsis thaliana. In the first setup, root hairs grow against an elastic obstacle until buckling. By measuring the critical buckling force, we determine the surface modulus and estimate the Young’s modulus of the cell wall, which aligns with previous measurements. Using a 1D elasto-viscoplastic model of root hair growth, we assess the excess pressure beyond the yield threshold (the driver of tip growth) and estimate the axial stiffness of the root hair, reflecting its elastic resistance to compression. For the second protocol, we designed a setup where a single root hair grows against a cantilever with variable stiffness, a technique adapted from our earlier work on rigidity sensing by animal cells. This method provides an independent estimate of the root hair’s axial stiffness, confirming our initial findings and suggesting that this stiffness primarily involves tip compression and depends mainly on turgor pressure, at least within the low deformation regime explored.

## Introduction

Root hairs (RH) are cylindrical extensions that develop from differentiated cells of the root epidermis. These highly elongated tubular structures, approximately 10 µm in diameter and up to a few millimetres in length(Meisner and Karnok, 1991), play a crucial role in nutrient uptake by increasing the root-soil exchange surface area by up to twofold(Jungk, 2001). To effectively grow and invade soils that present mechanical resistance, such as obstacles like rocks or compacted soil, RH must overcome these challenges.

The invasion of soil by RH is facilitated by turgor pressure, the motor of polarized tip growth, and by the mechanical support provided by the cell wall (CW) which is the primary component of the RH structure. Due to its rigid structure, the CW maintains cell integrity under turgor pressure and protects cells from mechanical stresses. The spatio-temporal regulation of its mechanical properties is essential for cellular growth. Under isotropic turgor pressure, the regulated processes of secretion, synthesis, and modification of the cell wall at the tip lead to polar (oriented) growth of the root hair(Dumais et al., 2006).

The CW, which is primarily composed of cellulose, hemicellulose, and pectin is present on both the shank and tip of the cell and is crucial for tip growth. The RH CW shows randomly oriented cellulose microfibrils at the tip and a weak alignment of cellulose microfibrils along the longitudinal axis in the shank(Akkerman et al., 2012; Newcomb and Bonnett, 1965). Measuring the mechanical properties of the RH CW is thus vital for understanding the complex process of cell growth and soil invasion.

Many studies have been devoted to investigating tip growth in resistant media, but with two main strategies. On the one hand, the analysis of tip growth as a whole in media of well calibrated mechanical properties, mainly for pollen tubes and root hairs(Pereira et al., 2024; Reimann et al., 2020). On the other, measurements of specific mechanical parameters of different tip-growing species. In particular, measurements of the Young’s modulus of the cell wall have been reported in various tip-growing systems, such as pollen tubes, hyphae, and fission yeast, subjected to bending or buckling experiments(Couttenier et al., 2022; Minc et al., 2009; Nezhad et al., 2013). For root hair cells, local measurements of the Young’s modulus and stiffness have been conducted using Atomic Force Microscopy (AFM)(Hirano et al., 2018; Shibata et al., 2022), and we have previously developed a flexion setup to measure the surface modulus of the CW of a growing RH(Pereira et al., 2023a).

In this context, we present here two original mechanical setups and protocols to characterize the mechanical properties of single growing RH of Arabidopsis thaliana. In the first setup, RH are allowed to grow against an elastic obstacle until buckling occurs. By measuring the critical force at buckling, we determine the surface modulus and estimate the Young’s modulus of the CW, which aligns well with values obtained from the flexion protocol(Pereira et al., 2023a). We then use a 1D elasto-viscoplastic model of RH growth to retrieve the excess pressure beyond the yield threshold (*i*.*e*. the motor of tip growth) as well as an estimation of the axial stiffness of the RH, *i*.*e*. its elastic axial resistance to compression. For the second experimental protocol, we designed an original setup to grow a single RH against a cantilever of variable apparent stiffness, a technique adapted from a setup we initially developed for characterizing rigidity sensing by single animal cells(Bufi et al., 2015; Mitrossilis et al., 2010). This protocol allows us to get an independent estimation of the apparent axial stiffness of RH, confirming the first measurement and suggesting that this apparent RH stiffness mainly involve RH tip compression and depend thus primarily on turgor pressure, at least in the explored low deformation regime.

### Buckling under its own load: insight into root hair cell wall elasticity

To measure the Young’s modulus of the cell wall of a growing root hair, we relied on a buckling experiment combining a technique previously used on individual animal cells(Desprat et al., 2006; Mitrossilis et al., 2010) and a buckling protocol used for yeast(Minc et al., 2009).

*Arabidopsis* roots and root hairs were grown on ½ MS agar medium using a microfluidic-like system previously described(Pereira et al., 2023b). This device enables high-resolution imaging under a microscope. Prior to the experiment, parts of the agar were removed to allow the root hairs to grow freely in liquid 1/2MS medium, meanwhile the root itself remained mechanically stabilized in agar to minimize drift during force application. To apply forces to the growing root hair tip, we designed a custom flexible glass microplate that functions as a cantilever of calibrated bending stiffness *k*. While growing, the RH progressively deflects the cantilever, thereby self-imposing a force *F(t)* = *k δ(t)* exerted on its tip, with *δ(t)*, the cantilever deflection (Figure 1A-C). This deflection, and thus the force supported by the RH, increases over time until it reaches a maximum (Figure 1C). This maximum corresponds to the critical buckling force Fc. Like many tip-growing cell(Burri et al., 2018; Minc et al., 2009), the RH buckles under the force generated by its own growth against an obstacle, and buckling is visible in the last image of Figure 1B.

**Figure 1:**
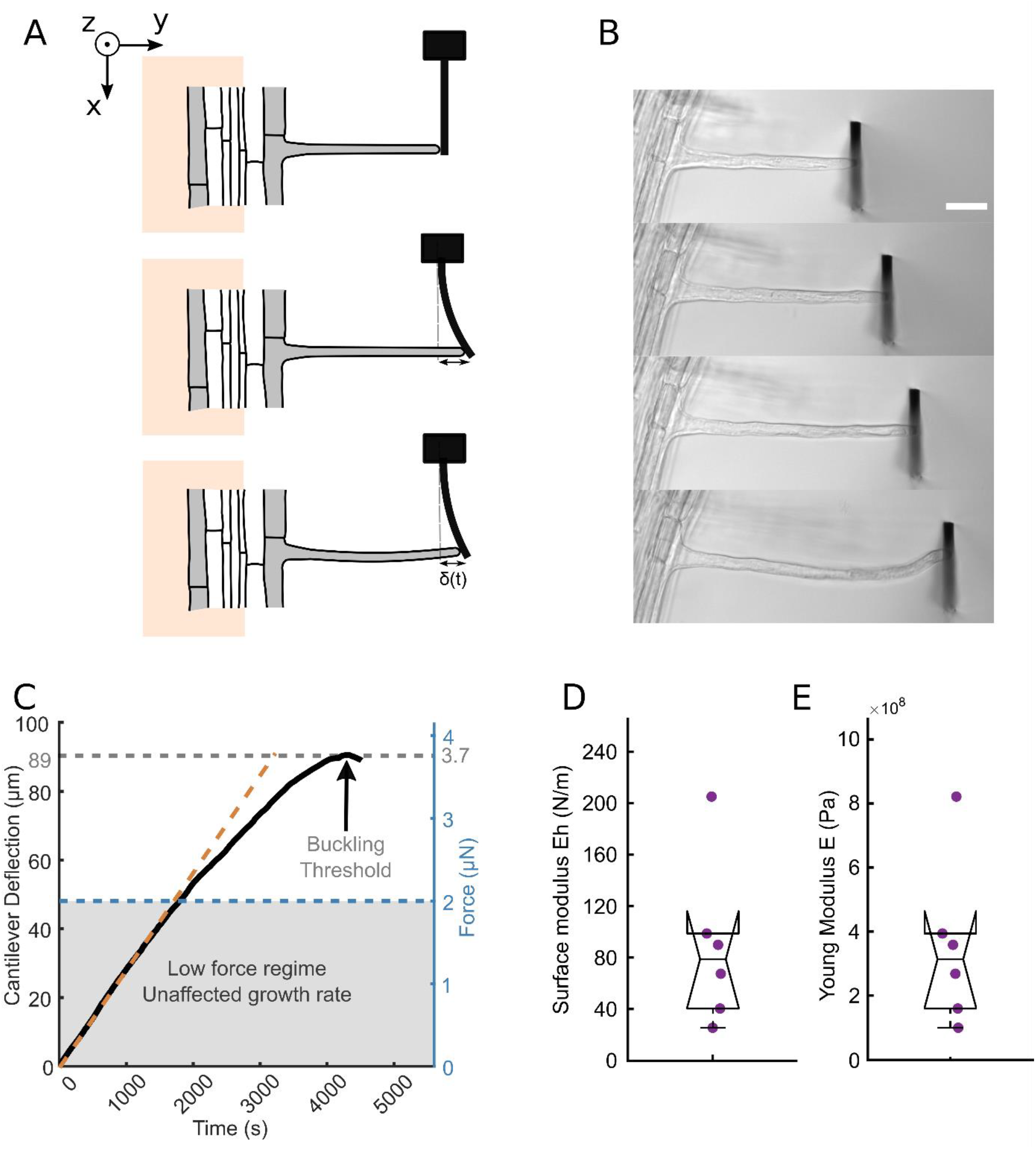
RH growth against a glass cantilever. **A**. Schematic representing the buckling experiment. The growing root hair progressively deflects the microplate and finally buckles. **B**. Time-lapse of a root hair growing against a microplate of stiffness *k* = 41.7 nN/µm (scale bar: 40 µm). **C**. Deflection of the microplate by a growing root hair over time. The corresponding level of generated force is represented on the right axis. The deflection reaches a maximum when the root hair buckles under the load applied by the microplate. The orange dotted line visualizes the linearity of the elongation curve in the low force regime, below 2µN. **D**. Boxplot showing the distribution of the measured surface modulus (n=6). **E**. Boxplot showing the distribution of the measured Young’s modulus (n=6).

For now, we use the critical buckling force Fc to estimate the surface modulus as well as the Young’s modulus of the cell wall. The root hair is considered as a hollow cylinder of radius *R*, length *L* and thickness *h*, with one end fixed (the RH basis clamped to the root) and the other end free to move laterally (the RH tip in contact with the glass plate). In these conditions, the critical force *F*_*c*_ is related to the Young modulus *E* and the thickness *h* of the cell wall by 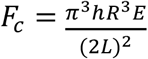. The RH radius and length are measured on the microscope images. We can then determine the cell wall surface modulus through the expression 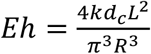. We measure *Eh* = 88 +/- 26 N/m (+/- SEM, n=6). To estimate the cell wall Young’s modulus, we assumed the cell wall’s thickness to be *h* = 250*nm*. This estimate is based on electron microscopy measurements of wild type root hairs. This gives a Young’s modulus *E* = 350 +/- 105 *MPa* (+/- S.E.M, n=6). These values are in good agreement with the values we previously measured on wild type RH through a bending experiment, with *Eh* = 113 +/- 24 N/m and *E* = 453 +/- 46 *MPa*(Pereira et al., 2023a). This agreement between values obtained by two independent methods validates both the values obtained for Eh and E for the RH wall, as well as the method of measuring Eh through the self-buckling experiment. Interestingly, the value of E we measure for single RH is close to the Young’s modulus of wild-type pollen tube measured at 350 MPa using fluid flow to bend the cell(Nezhad et al., 2013), as well as to that of fission yeast estimated at 101 MPa through buckling(Minc et al., 2009).

### Growth against an elastic obstacle: estimating growth pressure and RH stiffness

As long as buckling has not occurred, the deflection of the cantilever is equal to the elongation of the growing RH, and therefore its derivative (the slope of the curve of Figure 1C) gives the elongation rate of the RH. Here, we consider a 1D elasto-viscoplastic model to analyse the mechanics of RH growth against an elastic obstacle to get insight into the mechanical properties of the RH.

As detailed in(Dumais, 2021), such a model decomposes the root hair deformation during growth into an elastic (reversible) and a viscoplastic (irreversible) components. Indeed, form the mechanical point of view, the RH can be seen as an elastic cylinder with a yield fluid cap (Figure 2). For a freely growing RH, its current length can be expressed as:

**Figure 2:**
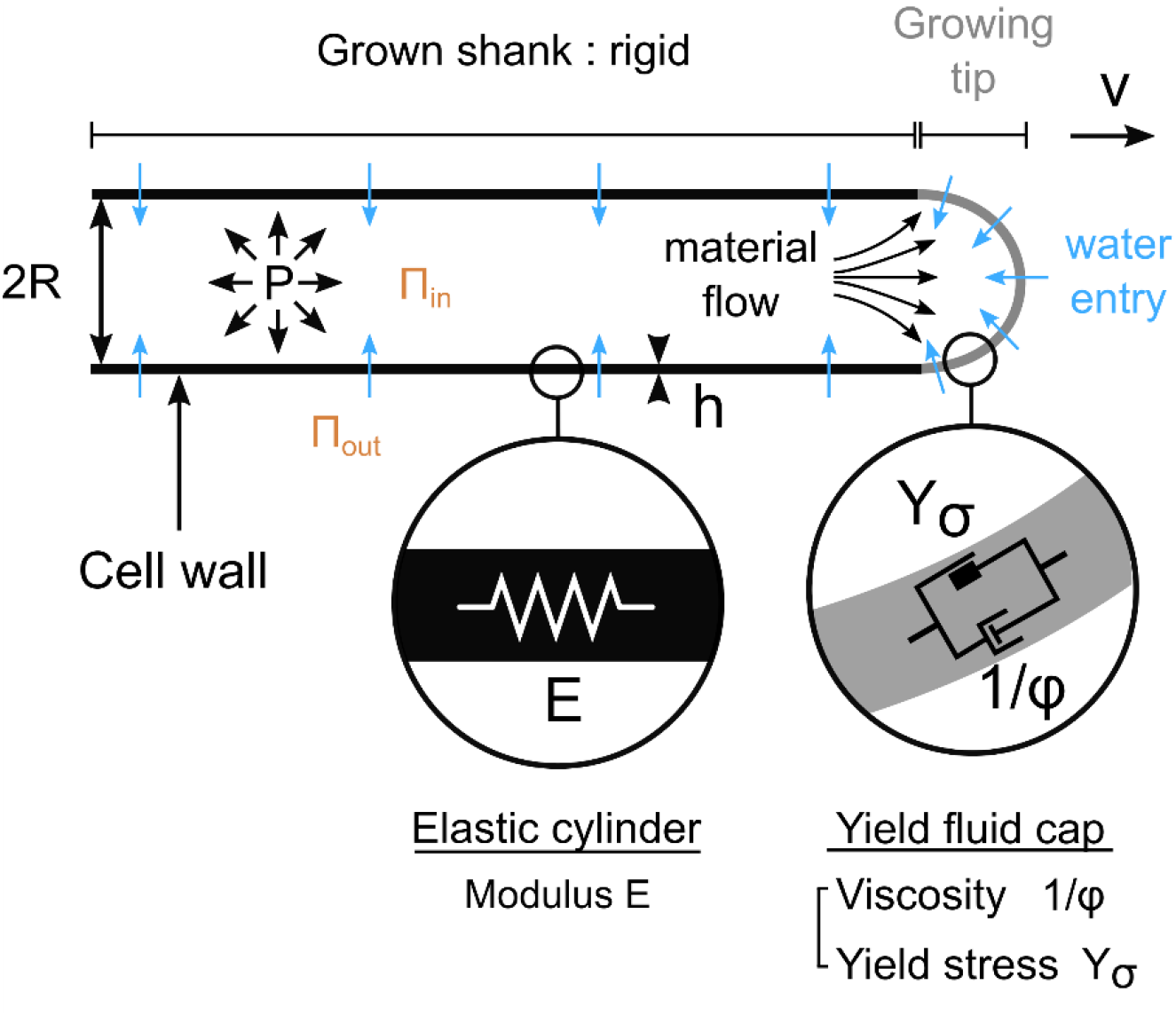
Root hair growth model. Schematic representing a root hair (RH) with a focus on the structural and mechanical parameters involved in the elasto-viscoplastic model describing its elongation and growth. The RH is composed of a pressurized tube with a shank surrounded by an elastic cell wall. At the tip the cell wall is described as a viscoplastic material. The turgor pressure P drives the growth at a speed v and only the tip is elongating. The root hair radius is noted R and the thickness of the cell wall is denoted h. Y and ϕ represent the physical parameters of the viscoplastic growth model. E represents the cell wall Young modulus. Water flows inside the RH because of an imbalance of water potential.

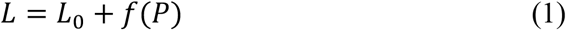

*L*_0_ corresponds to the plasmolyzed root hair cell length, i.e. when the turgor pressure *P* = *P*_*in*_ − *P*_*out*_ is set to zero (deflated RH). Thus, *f*(*P*) represents the change in length of the RH when it is put under pressure.

Irreversible growth corresponds to increasing *L*_0_ by adding new material to the cell wall at the tip of the RH. This is the true growth in the context of biology. Indeed, the reversible deformation stands for the elastic elongation due to pressure increase, such a growth component in steady state would imply a continuous increase of *P*. Considering turgor pressure as constant during free steady state RH growth, 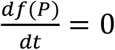 and the rate of RH elongation writes:

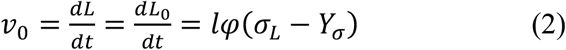

Where *l, φ, σ*_*L*_ and *Y*_*σ*_ are the typical length of the growth zone (∼ the tip), the extensibility of the cell wall in this zone (inverse of a viscosity), the longitudinal stress in the wall (along the RH axis), and the yield stress over which the tip flows, respectively. Equation (2) is simply the stress-strain relationship of a yield stress liquid.

When growing against an elastic obstacle like the glass microplate of Figure 1B, the RH is submitted to an opposing, compressive, force *F(t)* = *k*_*ext*_*δ(t)*T, with *k*_*ext*_ the stiffness and *δ*(*t*) the deflection of this external spring. The RH can then be seen as a whole as a spring with an effective stiffness *k*_*cell*_, representing the axial deformation under axial loading. In these conditions, the RH length is reduced by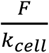 due to external compression:

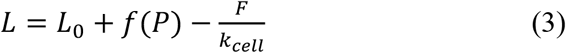

And, for constant *P*, the elongation becomes:

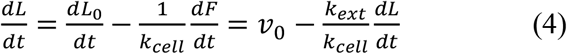

where we took into account the fact that, before buckling, cantilever deflection is equal to the cell elongation, *dδ* = *dL(t)*.

Equation (4) leads to:

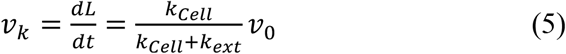

Implying a linear RH elongation over time, but with a reduced speed due to elastic compression, which involves a direct comparison of *k*_*cell*_ to *k*_*ext*_. We will use this expression later in the section dedicated to external stiffness modulation. For now, it must be noticed that (5) is only valid for a low force regime where the irreversible growth, i.e. *v*_0_, is unaffected by the compressive force. This regime is indeed observed on Figure 1C for forces less than 2 µN.

The force *F* externally applied on the RH reduces the longitudinal stress in the cell wall(Minc et al., 2009), reducing thus the growth rate:

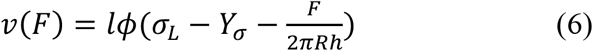

With R and h being the radius of the RH and its wall thickness respectively (Fig. 2). This expression can also be written as:

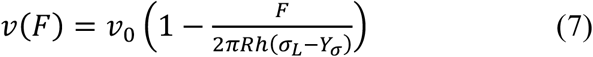

And the low force regime corresponds in fact to forces inducing small longitudinal wall stress 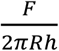 as compared to the excess stress (*σ* _*L*_ −*Y*_*σ*_) driving RH growth. Using 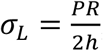 one can express *v*(*F*) as function of the excess (growth) pressure *P* _*G*_ = *P* − *Y* _*p*_ with 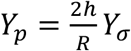 the yield pressure:

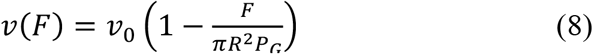

Replacing *ν*_0_ by *v*(*F*) in (4) one gets the differential equation controlling RH elongation over time and taking into account both the elastic compression and growth reduction effects imposed the cantilever facing the RH:

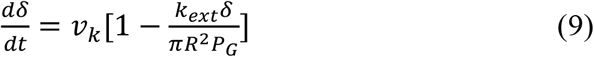

Written here in terms of the cantilever deflection *δ*(*t*) = *L*(*t*) − *L*_*i*_, with *L*_*i*_ the initial RH length, when contacting the cantilever. This leads to an exponentially saturating behaviour for RH elongation and force, at least before the onset of buckling:

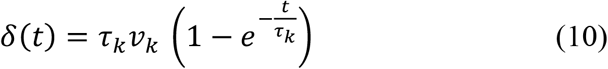

With

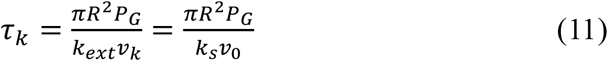

and

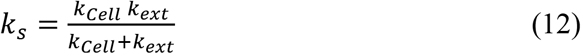

the apparent stiffness of the RH and the cantilever in series.

Fitting the deflection curves with Equation (10) we could get *τ*_*k*_ and *ν*_*k*_, and subsequently the values of the excess growth pressure *P*_*G*_ as well as the RH stiffness *k*_*cell*_ through:

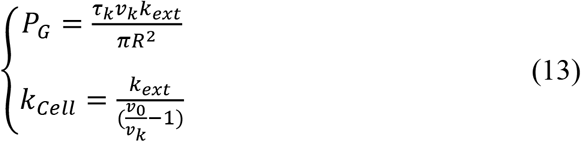

For the 6 cells tested, we found *P*_*G*_ = (1.07 +/- 0.23) 10^5^ Pa ≈ 1 bar, an estimation obtained in a non-invasive way for single growing RH.

To estimate *k*_*cell*_ we first measured *ν*_0_ for each cell as the tip speed before the RH enters in contact with the cantilever and then used Equation (13) to get the apparent RH stiffness. We found *k*_*cell*_ = 0.9 +/- 0.3 N/m ≈ 1 N/m. One can compare this value to a rough estimation considering that *k*_*cell*_ is mainly due to the stiffness *k*_*E*_ of the cylindrical RH shank (Supplementary Material) of which we have measured the surface modulus through RH buckling:

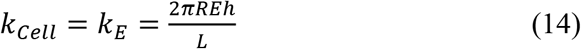

With *L*∼*1*00 µ*m, R*∼5 µ*m, Eh*∼100 *N*/*m*, one gets *k*_*E*_∼30 *N*/*m* which is an order of magnitude higher than the measured *k*_*cell*_.

At this stage, we sought to confirm the value of *k*_*cell*_ using another protocol. Indeed, as observed in Figure 1C, the cells buckle well before the force reaches its exponential saturation. The parameters of the adjustments, particularly the characteristic time *τ*_*k*_, are therefore poorly defined, which could lead to significant uncertainties in the estimation of *k*_*cell*_. Additionally, we assumed constant pressure, which may be questionable when monitoring growth over periods of about 20-30 minutes. Finally, the measured value of *ν*_0_ before contact between the RH and the cantilever, which reflects physical characteristics of the RH such as viscosity or yield stress, could also evolve over such long observation times.

To address these questions, we decided to conduct experiments on RH growth under conditions of time-varying external stiffness *k*_*ext*_. These experiments are detailed in the following section. However, it is worth noting that growth measurements are only conducted for a total of 40 seconds, and changes in stiffness are made in less than 0.1 seconds, thereby limiting any potential changes in *P* and *v*_0_ during the measurement. As we will see later, the measurements are taken in the low-force regime during which growth is linear over time and Equation (5) is valid. Using two different values of *k*_*ext*_ then allows us to even bypass the measurement of *v*_0_, making the measurements of *k*_*cell*_ free from the difficulties posed by the measurements through exponential growth curve fitting.

### Instantaneous growth modulation upon obstacle effective stiffness variation

To modulate the effective stiffness of the obstacle resisting RH growth, we adapted a technique we have originally designed to characterize rigidity sensing by single animal cells(Bufi et al., 2015; Mitrossilis et al., 2010). In practice, a glass microplate of calibrated stiffness is put in contact with a growing root hair as described in the previous section, but here we use a dual feedback loop to vary at will the force-deflection relationship, *i*.*e*. the apparent cantilever stiffness, thus changing the effective external stiffness *k*_*ext*_ of the obstacle facing the RH in less than 0.1 second (Figure 3).

**Figure 3:**
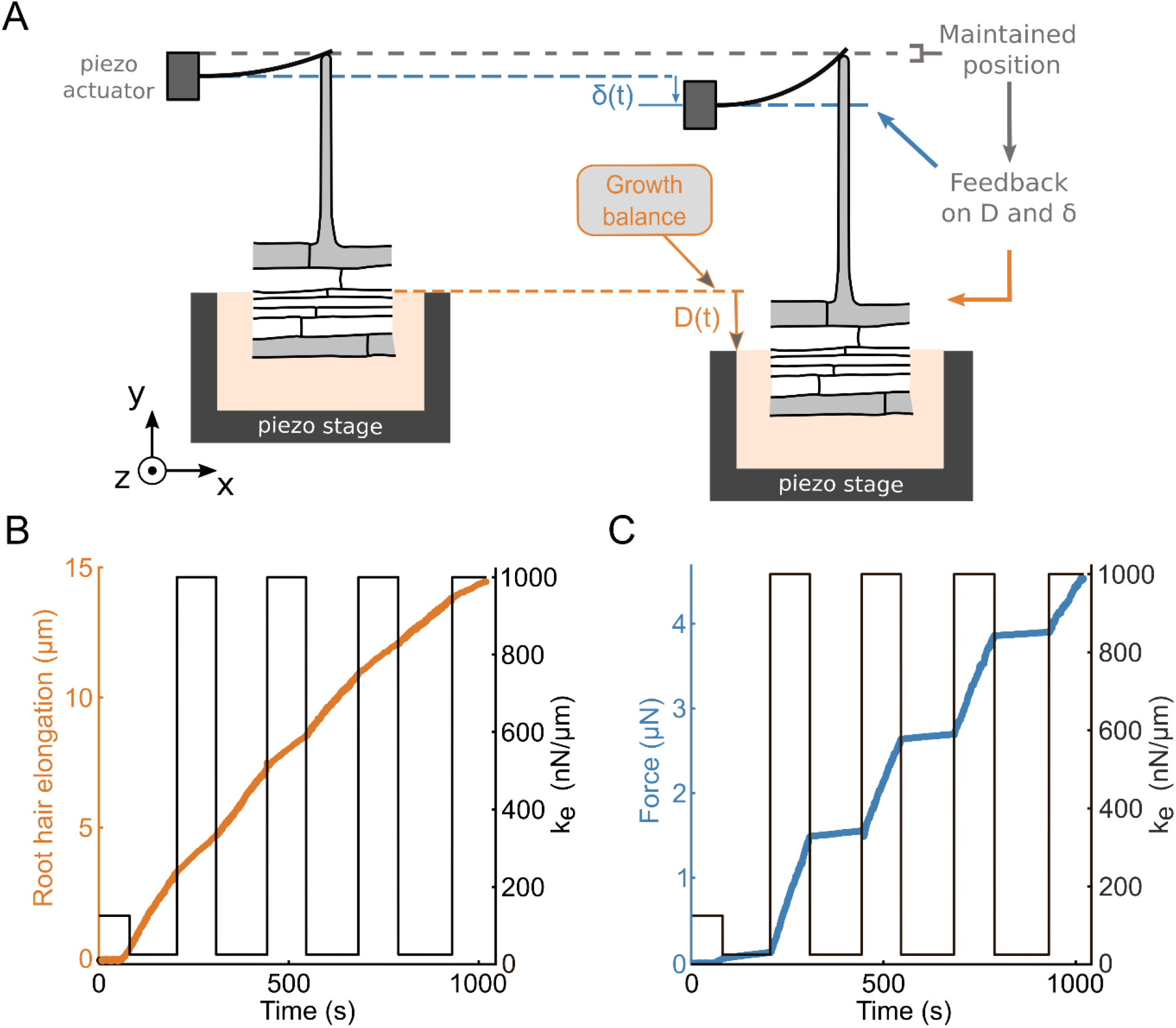
Instantaneous variations of the rates of elongation upon successive effective rigidity modulation. **A**. Schematic of an iterative step of the effective rigidity setup. The position of the contact between the RH and the microplate is maintained constant over time by double feedback signal: D(t) controls RH elongation, and *δ*(t) the deflection of the cantilever (a glass microplate of calibrated stiffness), and thus the force resisting RH elongation. **B-C**. RH elongation and force during successive changes in effective stiffness. **B**. Evolution of the RH elongation (orange) when the cantilever effective stiffness (black) is modulated over time. **C**. Evolution of the force (blue) in response to the same cantilever effective stiffness modulation (black) over time.

The plant is mounted on a sample holder on a piezo stage which allows to control the position of the RH base (Figure 3A). The microplate is also mounted on a 3D micromanipulator arm on a 3D piezo actuator to control the microplate base position. A dual feedback loop is then used to control both the positions of the cantilever and the plant. The position of the contact point between the root hair apex and the microplate is measured using a position-sensitive detector. This position is maintained constant by the dual feedback despite RH elongation during growth. To do so, the double feedback loop applies simultaneously a displacement *D*(*t*) to the root holder and a displacement *δ*(*t*) to the base of the cantilever. *D*(*t*) corresponds to a displacement of the root, and thus of the base of the root hair. Since the position of the contact between the RH and cantilever tips is kept constant during the experiment, *D*(*t*) is a measure of RH elongation. Symmetrically, *δ*(*t*), corresponds to the cantilever deflection. The effective cantilever stiffness *k*_*ext*_ is then given by the ratio between the increase in force *kδ*(*t*) and the increase in RH length *D*(*t*). Thus 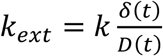 where *k* is the physical stiffness of the cantilever. The experimenter can adjust the effective stiffness, in real time, by modifying the ratio between *δ*(*t*) and *D*(*t*)(Mitrossilis et al., 2010). The figures 3B and 3C illustrate the response of a single RH to successive changes in *k*_*ext*_ between 25 and 1000 nN/µm. The elongation rate of the RH and the rate of increase of the force change instantaneously with each change in *k*_*ext*_. It is also noted that the elongation curve still overall resembles that of a saturating exponential as the force level increases, with a slope that decreases and thus makes it increasingly difficult to distinguish the jumps in speed. For the measurements of *k*_*cell*_, we therefore limited ourselves to studying the first jump, in the low-force regime, just after contact between the RH and the cantilever.

At the beginning of the experiments, a growing root hair was placed in front of a microplate, at a distance of a few micrometres, and brightfield images were taken every 2 seconds using a 40X objective to monitor growth. Initially, the effective stiffness was set equal to the actual physical stiffness of the microplate (*k* = 125 *nN*⁄µ*m*) by applying the same displacement to both the root and the microplate base (*δ*(*t*) = *D*(*t*)). Once the RH tip has reached the microplate and contact was established, the effective stiffness was set to 25 *nN*⁄µ*m* (Figure 4A). The root hair was then allowed to grow under this low stiffness condition for a few minutes, and the effective stiffness was then increased to either 500 or 1000 nN/µm depending on the experiment. Figure 4B shows an example of a sudden change in RH growth rate following an increase in effective stiffness from 25 to 1000 *nN*⁄µ*m*. To quantify this change, linear fits were performed on root base displacement data for 20 seconds before and after the stiffness change. For instance, in this particular experiment, the slopes shows a growth speed reduction from *ν*_25_ = *1*.*3*2 µ*m*⁄*min* at *k*_*ext*_ = 25 *nN*⁄µ*m* to *v*_1000_ = 0.79 µ*m*⁄*min* at *k*_*ext*_ = *1*000 *nN*⁄µ*m*. All curves were analysed for force levels < 0.4 µN (the force at stiffness changes was always < 0.25 µN), *i*.*e*. low as compared to the typical ∼2 µN axial force necessary to significantly reduce RH growth speed (Figure 1C).

**Figure 4:**
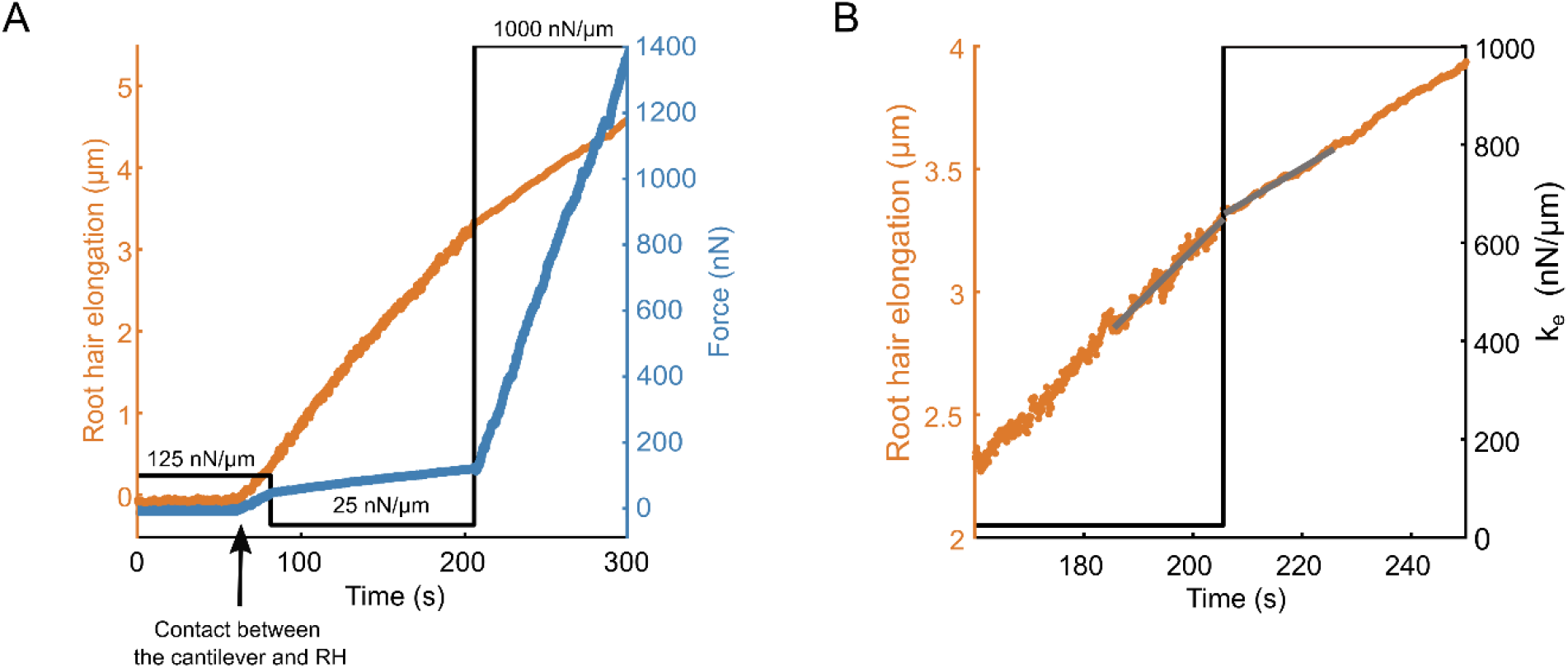
Instantaneous variation of RH growth rate upon effective stiffness modulation. **A**. Root hair elongation (orange) and force evolution (blue) over time when submitted to sudden jumps in external stiffness (black). Three successive effective stiffness values are explored successively: 125 nN/µm, 25 nN/µm and 1000 nN/µm. Until around 80 seconds, the RH is not in contact with the microplate, and no growth is measured. **B**. Zoom on the evolution of the elongation rate around a change in effective stiffness from 25 nN/µm to 1000 nN/µm (black). The grey segments are linear fits during 20 seconds before and 20 seconds after the change in stiffness.

To assess further le link between effective stiffness and growth rate, after the initial increase, the root anchoring in agar was checked by comparing the displacement controlled by the feedback loop and the observed root displacement on brightfield images (Figure S1).

Under these conditions, we consistently observed a decrease in the RH growth rate with increasing stiffness, and the decrease was greater with larger stiffness jumps. Specifically, jumps from 25 to 500 nN/µm resulted in a relative speed drop of 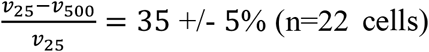 (n=22cells), while jumps from 25 to 1000 nN/µm caused a larger drop 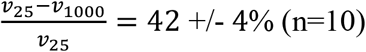 (n=10). This is in agreement with Equation (5) since it fits well *ν*_*k*_ data as function of the three *k*_*ext*_ values tested (Figure 5), although *ν*_*k*_ displays an important variability from cell to cell.

**Figure 5:**
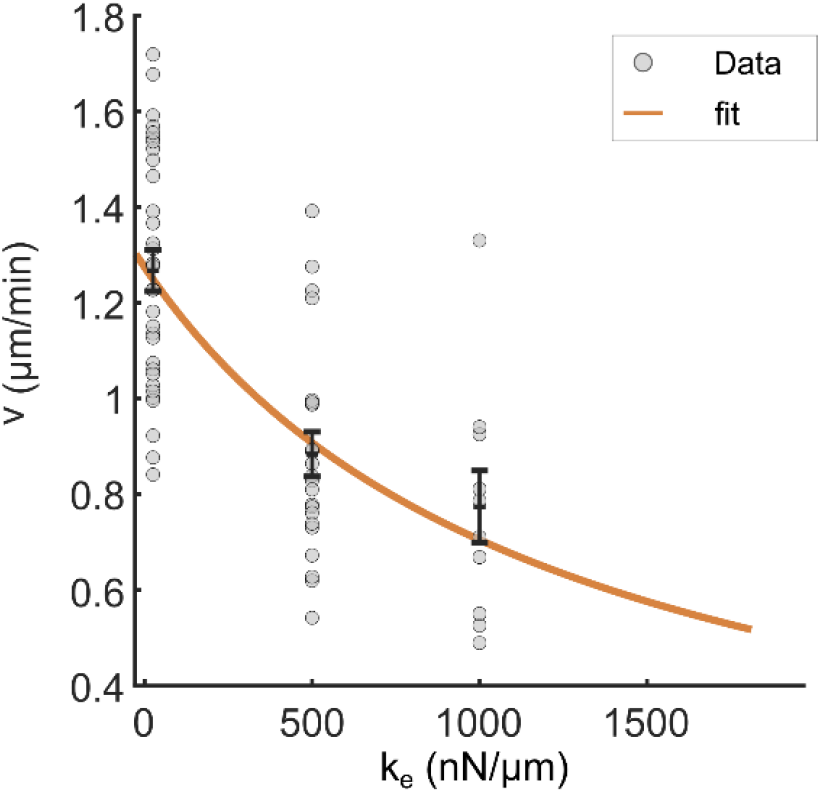
Elongation speed as a function of the cantilever effective stiffness for the cell population tested. RH growth speed represented as a function of the cantilever effective rigidity. The black error bar represents the mean +/- S.E.M for each rigidity (25 nN/µm (n=32), 500 nN/µm (n=22), 1000 nN/µm (n=10)). The orange curve represents a fit of Equation (5) to the data.

### Measuring RH stiffness from jumps in obstacle stiffness

Since measurements were taken in the low-force regime to ensure that the RH growth rate is solely a function of *k*_*ex*t_ and not of the force *F*, Equation (5) is valid, and we can express it for two different values *k*_*1*_ and *k*_2_ of *k*_*ext*_:

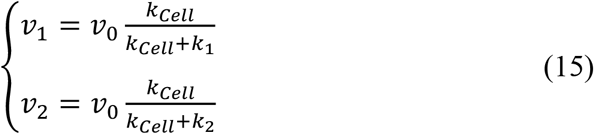

The ratio 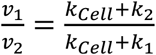 of the elongation rates allows one to express the RH stiffness *k* _*cell*_ as a function of the applied effective stiffness *k*_*1*_, *k*_2_ and of the measured speeds stiffness *ν*_*1*_, *ν*_2_:

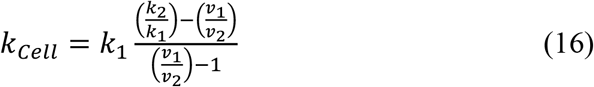

From the 25 to 500 nN/µm stiffness jumps, we found *k*_*cell*_ = 2.24 +/- 0.52 N/m (n=22 cells), resulted in a relative speed drop of 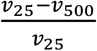, while jumps from 25 to 1000 nN/µm led to *k*_*cell*_= *1*.62 +/- 0.44 N/m (n=10). Pooling all the data leads to *k*_*cell*_ = 2.04 +/- 0.38 N/m (n=32 cells). Thus, we end up with *k*_*cell*_∼2 *N*/*m* confirming, by independent experiments and protocols, the ∼1 *N*/*m* found previously, and still an order of magnitude lower than the ∼30 *N*/*m* estimated for the stiffness of the RH shank.

## Discussion

Root hairs are cylindrical outgrowth of root epidermal cells that increases the surface of exchange with the soil and help the root penetrating the soil. Root hair growth is a typical example of apical (or tip) growth. This fundamental process allows plants, yeasts, and hyphae to invade their environment. This process thus involves complex mechanical aspects and numerous associated parameters, such as turgor pressure, the elastic modulus of the cell wall, the effective viscosity, and the yield threshold of the growth zone. To enhance our understanding of apical growth and validate realistic models, it is necessary to develop experimental devices and protocols to measure these parameters in vivo.

Here, we have described a micromanipulation device and original experimental protocols for measuring mechanical parameters characterizing the apical growth of root hairs when they encounter obstacles, a situation that is quite typical for this type of cellular extensions which necessarily involves soil penetration. First, through a self-buckling experiment, we were able to measure the surface modulus (*Eh* ∼ 100 N/m) and then estimate the Young’s modulus of the root hair wall (*E* ∼ 400 MPa). Then, by analysing the growth law of the root hairs over time, we measured the excess pressure above the yield threshold (*P*_*G*_ ∼ 1 bar), as well as the apparent axial stiffness of the root hairs (*k*_*cell*_ ∼ 1 µN/µm), which presumably allows them to limit their compression during soil penetration. Since the value found for cellular stiffness was an order of magnitude smaller than expected, based on the surface modulus value of the wall (*k*_*E*_ ∼ 30 µN/µm), we performed an independent new measurement using a variable stiffness device that we specifically developed. We then confirmed the low value of the axial stiffness of the root hairs (*k*_*cell*_ ∼ 2 µN/µm). These results highlight the importance of taking into account elastic deformations (*i*.*e*. reversible deformations) in mechanically constrained growth experiments. Indeed, if the applied force varies in time, the observed elongation rate is not equal to the growth rate due to cell wall yielding.

This result, which is a priori surprising, could be explained by a low apparent stiffness of the growing tip of the RH, which is in fact the part directly in contact with the obstacle. The wall in this growth zone is considered as a liquid sheet flowing under the effect of the surface stress generated by the difference in hydrostatic pressure *P* on either side of the wall. Pressing on the tip of the RH therefore amounts to increasing the contact surface area between the obstacle and the RH, and thus the pressure forces. The tip of the RH would then behave like a spring with stiffness *k*_*p*_ = π*RP* (at least for small indentations, see Supplementary Material and Durand-Smet et al(Durand-Smet et al., 2017)). Given that the radius of the RH is typically 4-5 µm, the measured stiffness of the RH *k*_*cell*_ of approximately 2 µN/µm would therefore imply a pressure *P* of about 1.5 bar, which is quite plausible and certainly in the right order of magnitude.

At this stage, it is tempting to combine the values of the excess pressure *P*_*G*_ (obtained via exponential growth) and the hydrostatic pressure *P* (estimated here assuming *k*_*cell*_ ∼ *k*_*P*_) to estimate the pressure growth threshold *Y*_*p*_ = *P* − *P*_*G*_. This yields *Y*_*p*_ = 0.5 bar, a quantity for which there is currently no measurement and thus no reference in the literature. However, caution is necessary, particularly because *P*_*G*_ and *P* were estimated from two different cell populations, and the data in Figure 5 clearly show that cellular variability is significant. For example, growth rates at very low external stiffness (and thus close to the free growth rate *v*_0_) vary between 0.8 and 1.8 µm/min. Indeed, the average speed for the sample used to estimate *P* is approximately 1.3 µm/min, whereas it is about 1.8 µm/min for the cells used to measure *P*_*G*_. To address these challenges, we are currently working on developing techniques and experimental protocols that would allow us to obtain specific and independent measurements of turgor pressure *P* on one hand, and the yield pressure *Y*_*p*_ of the wall in the growth zone on the other. Furthermore, to validate the idea that the tip of the RH acts as a spring with stiffness *k*_*P*_, which is 10 times smaller than the typical stiffness of the rest of the cell *k*_*E*_, it would be necessary to develop precise measurements of the deformation of the cell wall along the entire contour of the RH.

## Material and methods

### Device preparation

The microfluidic-like system(Pereira et al., 2023b) used to grow the roots was fabricated using a custom-made PVC mold featuring a single channel measuring 270 µm in height and 1 cm in width. A 10:1 mixture of PDMS base and curing agent (Sylgard 184, Dow Corning) was poured into the mold and cured overnight at 65 °C. The cured PDMS chip was then bonded to a glass microscope slide using a plasma cleaner (Harrick Plasma, PDC-002-CE). Then the channel was filled with ½ MS (Murashige and Skoog) medium supplemented with 5% sucrose (w/w) and 1% agar (w/w) (Duchefa, plant agar), with the pH adjusted to 5.7. Finally, a 0.5 cm thick layer of the same medium was poured around the PDMS chip.

### Plant culture

We used *Arabidopsis thaliana* seeds expressing the fluorescent nuclear envelope marker pSUN1:SUN1-GFP(Graumann et al., 2010). Seeds were sterilized and then stratified for 48 hours at 5 °C in 1 mL of ½ MS (Murashige and Skoog) medium supplemented with 5% sucrose (w/w). After stratification, the seeds were planted near the entrance of the channel on the agar-filled microfluidic device. The system was then enclosed in a Petri dish sealed with microporous tape and placed at a 45° angle relative to the vertical inside an incubator (Sanyo MLR-351H). Growth conditions consisted of a 16-hour light phase at 20.5 °C and an 8-hour dark phase at 17 °C, with 65% humidity.

### Cantilever preparation and calibration

The microplates were made by stretching (Narishige PB-7 puller, Japan) a glass plate, then cutting it and fusing it with a glass capillary as described in a previous study(Desprat et al., 2006). The calibration of the glass microplate consisted in calibrating it with a standard microplate as described before(Desprat et al., 2006). The microplates used in buckling experiments are placed along the vertical axis whereas in the stiffness experiments the microplate is placed parallel to the imaged plan to ensure a correct detection of its position by the position sensitive detector.

### Force application and data collection

Prior to the experiment the plant and the microfluidic-like system were placed on a microscope holder. Using tweezers and a scalpel, the agar on the side of the root tip was removed and replaced with ½ MS (MS Murashige and Skoog) liquid medium supplemented with 5% sucrose (w/w), PH 5.7. The system was then observed either with a 20×0.45 NA or a 40 × 0.55 NA objective under an IX83 Olympus microscope. The root was positioned at an angle to ensure that a single root hair grew in contact with the microplate. During the buckling experiment RH growth and the cantilever deflection were measured using brightfield images. In the variable stiffness set-up, the microplate was placed in position using a 3D micromanipulator system, ensuring that its position could be recorded by a position-sensitive detector. (S3979 one-dimensional Position Sensitive Detector Hamamatsu, France). The feedback loop allowing to modulate cantilever stiffness was adapted from a previously described experiment(Mitrossilis et al., 2010). The positions of the piezoelectric stage and piezo actuator controlling the cantilever base position were recorded during the experiment. Bright field images were taken to assess the root hair growth direction. In all experiments, RH elongation speed was estimated by fitting the piezoelectric position versus time.

### Statistical analysis

The Data were analysed using Matlab and FIJI. The data are presented as mean values +/- s.e.m. For the values determined by a fit, the Matlab Curve Fitting tool was used and data are presented as fitted value +/-half-width of the 95% confidence interval.

## Data availability

Data will be made available upon reasonable request

## Author contributions

DP, TA and AA developed the conceptual framework and designed the study. AA sought funding. TA, DP and LC performed the experiments. TA performed the quantitative analysis of the results. DP, TA and AA analysed the findings and wrote the manuscript.

## Acknowledgements

We thank Marie-Edith Chabouté for providing SUN nuclear envelope marker lines. We thank Pauline Durand-Smet and Etienne Couturier for fruitful discussions.

