## Supplementary material for "Micro-mechanical approaches to characterize tip growth: Insights into Root Hair Elasto-Viscoplastic Properties": Suplementary material

### 1) Shank stiffness $k_E$

Outside the growth zone at the tip, the root hair can essentially be considered as a hollow cylinder with a radius R~5 µm, a length $L$, a thickness h~250 nm, and composed of an elastic material (the cell wall) with a Young Modulus E~400 MPa. When an axial force $F$ is applied on the section of the cylinder over an area ~$2\pi Rh$, it leads to a longitudinal stress $\sigma$:

$$\sigma = \frac{F}{2\pi Rh} = E\frac{\Delta L}{L}$$

With $\Delta L$ the subsequent change in cell length, and $\frac{\Delta L}{L}$ the corresponding strain. This relationship can be expressed in terms of force and elongation:

$$F = \frac{2\pi RhE}{L}\Delta L$$

Thus giving the shank stiffness:

$$k_E = \frac{2\pi REh}{L}$$

### 2) Tip stiffness $k_p$

At the tip of a growing root hair, the cell wall is flowing under the stress generated by the pressure jump $P$ across the wall. The tip can roughly be considered as an inflated soft hemisphere of radius R. When a flat glass microplate (the cantilever) indents the tip, the situation resembles that of Hertz contact between a sphere and a semi-infinite plane. The radius $a$ of the contact area is then given by:

$$a \sim \sqrt{R\delta}$$

With $\delta$ the indentation depth of the tip which is equal to the change in RH length $\Delta L = \delta$. The force applied by the cantilever can then be expressed as:

$$F = \pi a^2 P = \pi R\delta P = \pi RP\Delta L$$

And the apparent stiffness of the RH tip writes:

$$k_p = \frac{F}{\Delta L} = \pi RP$$

#### 3) Root anchoring

A poorly anchored root might be displaced when force is applied at the root hair tip, leading to errors in in the measured RH elongations. To check root anchoring, we compared the displacement applied by the feedback loop to the stage holding the root (black curve of Fig. S1) with the actual displacement of the root as observed in brightfield images (red). As shown in Figure S1, the two displacements overlap very well. The mismatch was not more than 500 nm over tens of microns displacements, confirming thus excellent root anchoring.

**Figure S1**

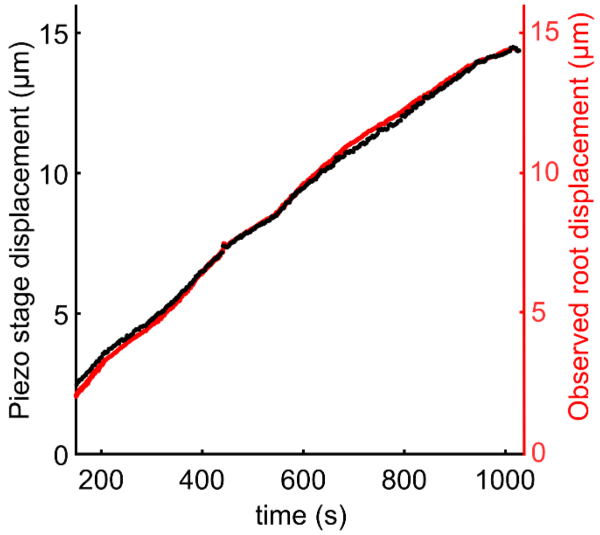


**Figure S1. Illustration of the good root anchorage.** During the variable stiffness experiment displayed in Figure 3, the stage displacement (black), imposed by the feedback loop on the piezo actuator controlling the root position, overlaps well with the actual root displacement (red) visualized through brightfield imaging and measured using the template matching tool of FIJI.

### 4) Correlation between feedback noise and effective stiffness:

A close look on the elongation and force curves during variable stiffness experiments reveal an uneven distribution of the noise on the signals. The RH elongation curve is noisier when the effective stiffness is low, whereas the force curve becomes noisier when stiffness is higher. This is because the feedback command is unevenly distributed among the stage displacement (RH elongation) and the cantilever base displacement (imposing deflexion and thus force). Indeed, the effective stiffness is set by the ratio between these two signals. At low effective stiffness ($k_{feedback} < k_{cantilever}$), a large fraction of the feedback command signal is devoted to elongation, whereas only a small fraction is devoted to force increase through cantilever deflection. In these conditions the feedback loop noise gets also unequally distributed between the signals. Thus, at low stiffness elongation measurements are noisier, whereas at high stiffness force measurements are noisier (Figure S2). This is particularly evident from the zoom on the elongation curve during stiffness change (Figure 4B). indeed, as visualized in Figure S2, the sum of the noises of the 2 signals is constant over time. This sum corresponds to the intrinsic noise of the feedback loop which is unevenly distributed between elongation and force depending on the chosen effective stiffness value.

**Figure S2**

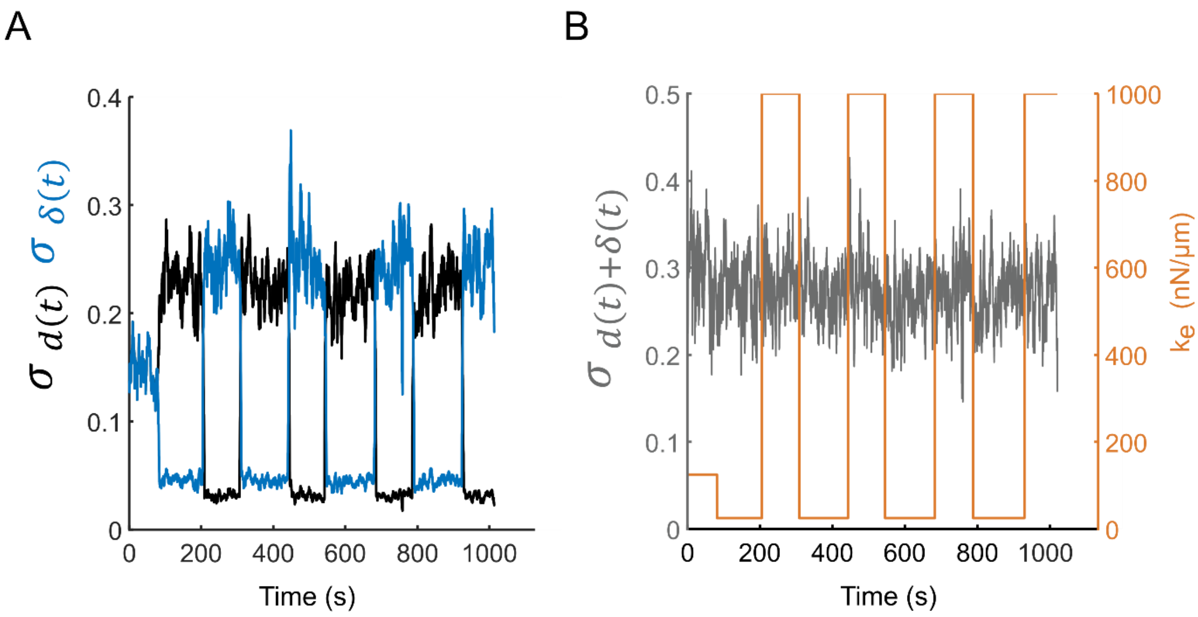


**Figure S2. Noise of the feedback loop.**

**A.** The black and blue curve display the standard deviation of the displacements $d(t)$ (elongation) and $\delta(t)$ (force) measured in a sliding window of 50 points (~5s) applied by the double feedback loop during the experiment showed in figure 3. **B.** The gray curve displays the standard deviation of the sum $d(t) + \delta(t)$ measured in a sliding window of 50 points. The orange curve represents the cantilever effective stiffness.
